# TraitProtNet: Deciphering the Genome for Trait Prediction with Interpretable Deep Learning

**DOI:** 10.1101/2024.03.28.587180

**Authors:** Sijun Wang

## Abstract

Genome data is far from fully explored. We present TraitProtNet, an innovative deep learning framework for predictive trait profiling in fungi, leveraging genome data and pretrained language models. The use of Integrated Gradients and bioinformatic analysis provides insights into the model’s interpretability, complementing traditional omics by highlighting the difference between protein importance and expression levels. This framework offers significant potential for future applications in both agriculture and medicine.

## Main

Advancements in sequencing technologies like Next-Generation Sequencing (NGS) and Mass Spectrometry (MS) have greatly enhanced accessibility to biological sequence data, with the NCBI database now housing 14,920 genomes for Eukaryota. Despite this abundance, the complexity of deciphering the intrinsic nature of nucleotide and protein sequences presents a formidable challenge, with traditional methods like BLAST (Altschul *et al*. 1990) often falling short. The evolving field of bioinformatics is increasingly leveraging machine learning (ML) and deep learning (DL) techniques, proven effective in handling the intricate patterns of biological sequences. For instance, AlphaFold2’s success in predicting protein structures signifies a leap in understanding protein functionalities encoded within amino acid sequences (Jumper *et al*. 2021). Protein language models (pLMs), adapted for biological sequences, are breaking new ground in identifying patterns and functions within protein sequences (Rives *et al*. 2021; Elnaggar *et al*. 2022).

The complexity of genomic sequence extends beyond the index encompassing functional information to include the spatial organization of chromosomes, gene regulation, and their interactions, underscoring the challenges in decoding genomes (Raser and O’Shea 2005; Fudenberg, Kelley and Pollard 2020). While substantial progress has been made in developing methods to interpret genomic data, developing a comprehensive tool that leverages this data to predict a wide array of traits across different species is still an unresolved challenge (Cao *et al*. 2022; Chen *et al*. 2022).

In this study, we introduce TaritProtNet, an innovative framework that integrates the pre-trained pLMs ESM-2 (Lin *et al*. 2023) with convolutional neural networks (Fig. 1A). First, we evaluated the capability of our acquired embeddings to represent proteins effectively. We analyzed the representations of homologous proteins as well as proteins with divergent sequences and structures. For each reference sequence, BDC60758, YP_010799217, and XP_009650171, we selected 100 sequences with high sequence similarity. Importantly, the three reference sequences were chosen based on their distinct functional attributes to prevent bias in the evaluation process.

**Fig. 1.**
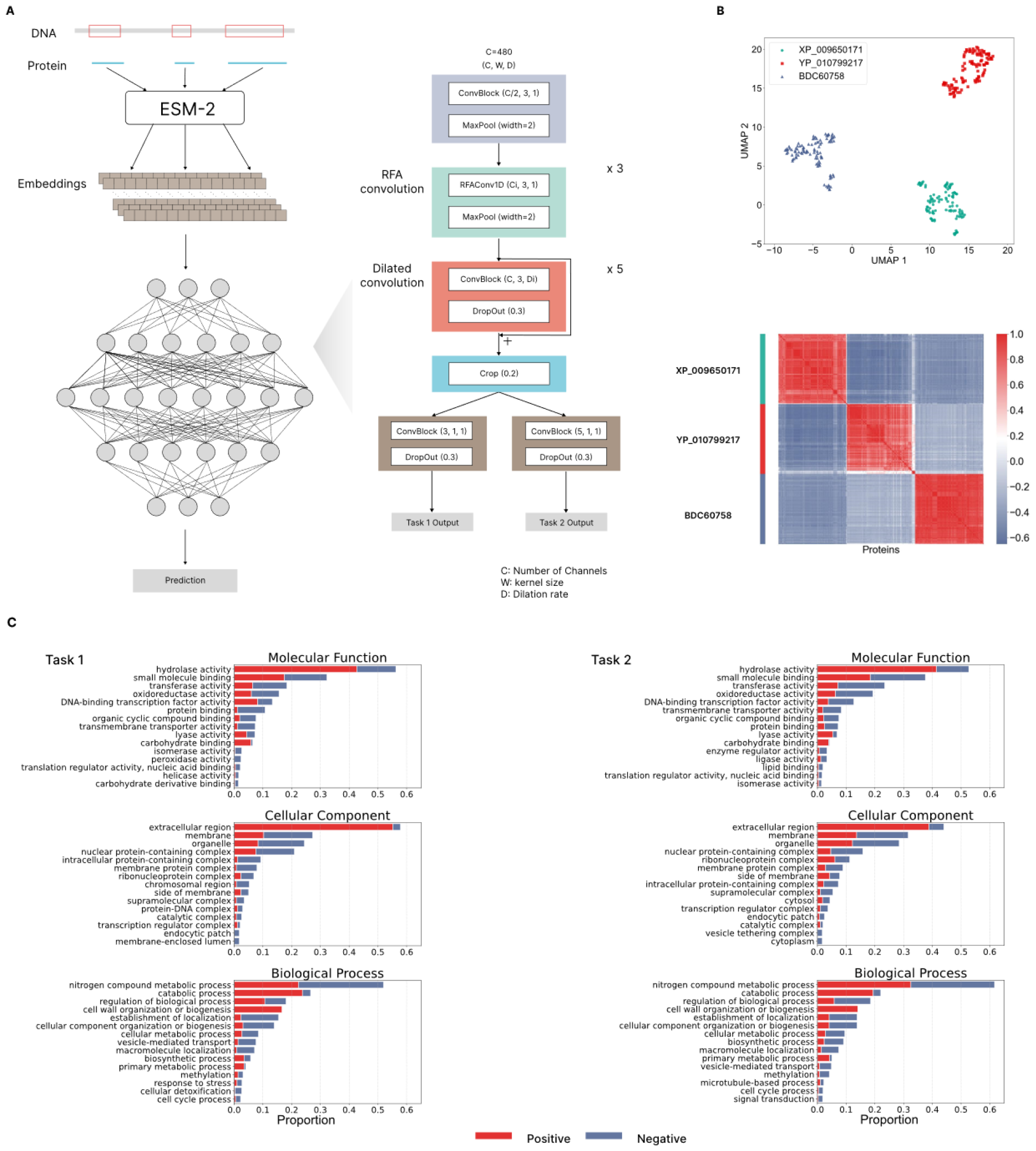
**A**, Schematic representation of the model architecture. **B**, UMAP visualization of protein sequence embeddings, with reference sequences XP_009650171, YP_010799217, and BDC60758, demonstrating clustering according to sequence similarity. The adjacent heatmap displays pairwise correlations between proteins, with red indicating a higher correlation. **C**, Bar charts of GO enrichment results for Tasks 1 and 2, illustrating the proportion of proteins associated with specific GO terms in Molecular Function, Cellular Component, and Biological Process categories, denoted by red for positive attribution and blue for negative attribution.

Uniform Manifold Approximation and Projection (UMAP) (McInnes, Healy and Melville 2020) are employed for dimensionality reduction to illustrate the discriminatory power of the embeddings generated by ESM-2 with 35 million parameters. The result revealed that the protein sequence embeddings aggregate into three distinct categories, each corresponding to a specific reference sequence, indicating the model’s capability of differentiating among proteins from disparate families (Fig. 1B). Furthermore, the heatmap of pairwise correlations among proteins corroborates these observations. The embeddings of proteins with sequence similarity to the same reference sequence exhibit higher correlation values (Fig. 1B).

Utilizing a dataset of 702 fungal genomes, we proposed a new data type built upon the embeddings, where each data point encompasses all proteins from a species. Despite the dataset’s relatively small size, this architecture demonstrates exceptional predictive performance. For a binary classification task to distinguish between different growth forms of fungi, TraitProtNet achieved an Area Under the Receiver Operating Characteristic curve (AUROC) of 0.992, closely aligned with the AUROC of 0.993 for BPNet (Avsec *et al*. 2021b). In predicting fungi’s primary lifestyle, the proposed model recorded an AUROC of 0.948 and a weighted F1 score of 0.837. These metrics are 0.062 and 0.197 points higher, respectively than those achieved by BPNet, indicating an improved performance in capturing the common features of fungi sharing same primary lifestyle.

Integrated Gradients (IG) (Sundararajan, Taly and Yan 2017) are used to calculate attribution scores for proteins, aiding their analysis with established bioinformatic methods. We focused on seven species with mycelium, a crucial fungal structure for life cycles, and identified as plant pathogens to interpret TraiProtNet harnessing the link between mycelium presence and pathogenicity (Xia *et al*. 2019). Gene Ontology (GO) enrichment analysis revealed that hydrolase activity, especially related to glycosidic bond modification or degradation, is a significant category of proteins with high attribution scores in both tasks, aligning with their pivotal roles in cell wall modification or degradation, a key process in pathogenic interactions and mycelium development (Fig. 1C) (Fang *et al*. 2019; Rafiei, Vélëz and Tzelepis 2021).

Furthermore, while DNA-binding transcription factors (TFs) activity was found in both tasks, they exerted a more pronounced positive influence in Task 1 (Fig. 1C and B). This observation highlights the capacity of our model to discern the subtle interactions within complex protein networks, tackling the existing challenges in differentiating the effects of TFs on fungal development and virulence, as well as establishing causality between these elements (John *et al*. 2021). In addition, Protein-Protein Interaction (PPI) analysis for the second task revealed a focus on proteins involved in plant cell degradation, crucial for pathogenic fungi’s nutrient acquisition (Kubicek, Starr and Glass 2014). PPI of another task emphasized transcription regulation proteins, indicating their potential importance in mycelium formation (Figs. 2A and B).

**Fig. 2.**
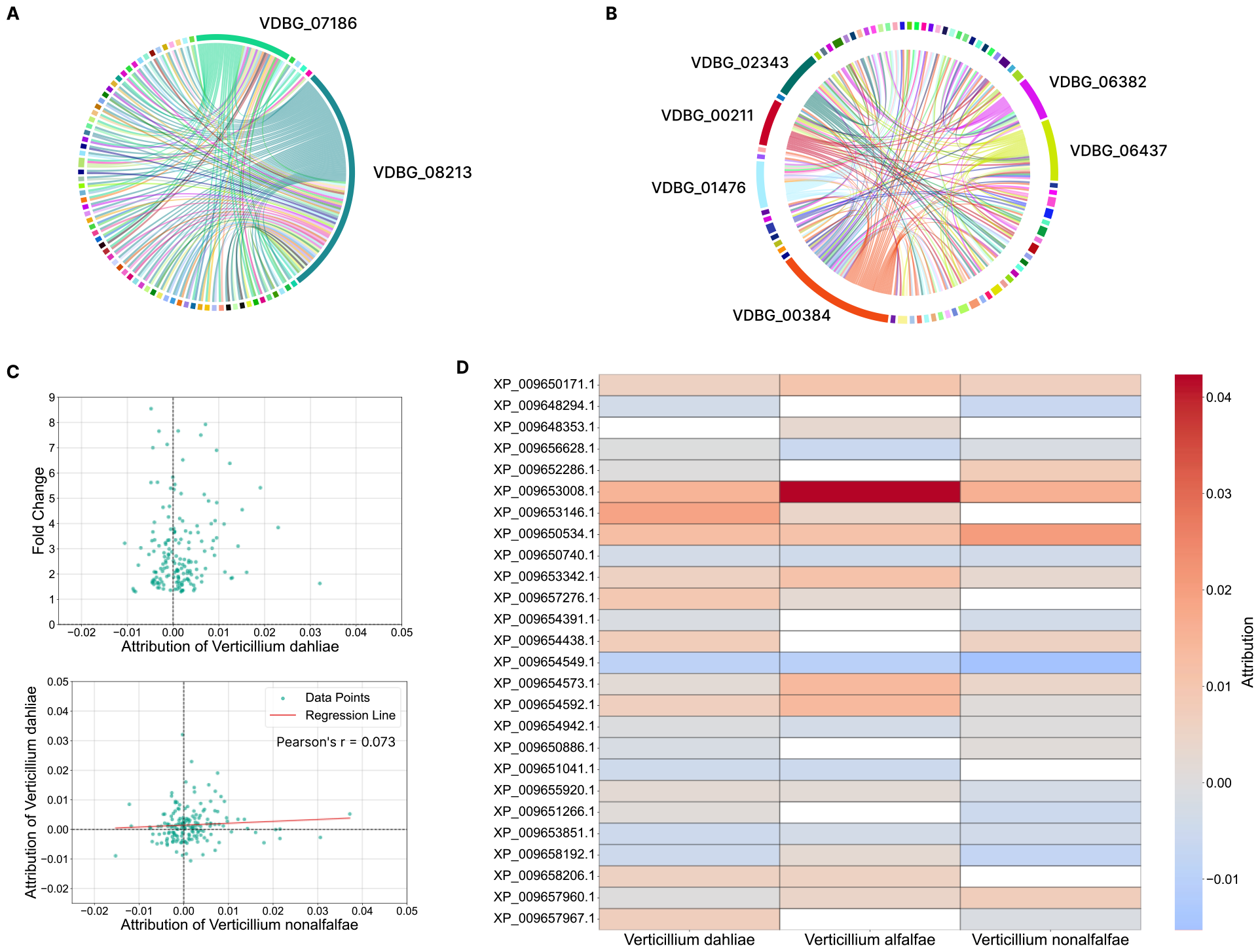
**A**, PPI network for high attribution proteins in *Verticillium alfalfae* for task 1 showcasing interactions with multiple proteins, with varying widths indicating the coverage of interactions. **B**, PPI network for high attribution proteins in Verticillium alfalfae for task 2 showcasing interactions with multiple proteins, with VDBG_00384 being the most active protein in the network. **C**, Scatter plot illustrating two distinct relationships: the first demonstrating the association between attribution scores and experimentally observed Fold Change in proteomic data for Verticillium dahliae, and the other displaying the non-linear correlation between model attribution scores across two different species. **D**, Attribution scores across three Verticillium species, with color intensity representing the magnitude of the attribution.

In our analysis of 181 upregulated proteins from experimental proteomic data of *Verticillium dahliae* (Wang *et al*. 2023), we observed no significant correlation between experimentally detected Fold Change and attribution values (Fig. 2C). This discrepancy may originate from the intrinsic limitations of IG and the model’s emphasis on identifying common patterns across species. However, it also calls for a reevaluation of the traditional omics approach that equates high expression levels with importance, which could introduce biases in identifying target proteins (Al-Amrani *et al*. 2021; Nakayasu *et al*. 2021). Despite the lack of correlation, and varying attribution scores among homologous proteins across species highlighted by a low Pearson’s correlation coefficient, 40 proteins consistently showed positive attribution scores in all three species (Fig. 2C). Furthermore, proteins with a Fold Change exceeding 5 presented a discernible pattern in attribution scores, despite variations, even for those lacking homologous counterparts in one or more of the species, suggesting a nuanced relationship between their true importance and their contributions to model predictions (Fig. 2D).

TraitProtNet emerges as a groundbreaking framework that leverages deep learning to unlock the vast potential of genomic data, refining protein sequence information through pretrained LMs for high-accuracy trait prediction. The adaptability of the proposed framework extends to applications in species classification and biosynthetic validation, with promising prospects for clinical feature detection in human genomes. The strong interpretability of TraitProtNet also paves the way for its use as a complementary tool in bioinformatics, shifting the focus from protein abundance to importance. This shift enhances the likelihood of identifying proteins with potential benefits for agriculture, medicine, and humanity at large, marking a significant advancement in the field.

For instance, by utilizing NGS to obtain the genomic data of a large population and simultaneously collecting their phenotypic traits to create a dataset, we can train our model on this information. This would enable us not only to predict individual traits, including the likelihood of disease or even simple characteristics like hair color at an early stage but also to trace back to the protein networks that contribute to these traits, potentially leading to targeted solutions. However, further validation is necessary to fully assess its ability to capture mutations at the amino acid level.

## Methods

### Data Preparation

The dataset utilized in this study originates from FungalTraits, which offers information on fungal species’ names along with 17 lifestyle-related traits of fungal and stramenopila genera (Põlme *et al*. 2020). We focused on growth and lifestyle traits, consolidating these into broader categories to mitigate data imbalance. Subsequently, we confirmed the availability of corresponding sequence data for these traits and retrieved them from NCBI.

### Acquiring Embeddings with ESM-2 and Handling Long Protein Sequences

Protein sequence embeddings were obtained using the ESM-2 model, which features 35 million parameters. To represent an entire protein sequence by a single vector, we averaged embeddings across the sequence length dimension. For sequences exceeding 1000 amino acids, we employed a sliding window approach with a 500 amino acid size to process the data, subsequently averaging these embeddings to obtain a comprehensive representation (Oliveira, Pedrini and Dias 2023). We then concatenated the vectors of all proteins belonging to a species to create a single data point.

### Homologous Protein Identification and Data Visualization

Homologous proteins were identified utilizing BLAST. Dimensionality reduction was performed using UMAP with the parameters: n_neighbors set to 10 and min_dist to 0.5.

### Dataset Preparation and Data Augmentation

We divided the dataset into a training set (70%) and a test set (30%), with half of the test set reserved for cross-validation and the entirety of the test set used for evaluation. To maintain the independence of the test dataset, we included species that do not share a genus with any species in the training data, ensuring a robust and unbiased evaluation of the model’s performance.

Given the limitations of a small dataset, we implemented data augmentation techniques for training data by randomly deleting 10% of vectors in a sequence and shuffling the order across the length dimension, thereby expanding the dataset to three times its original size.

### Architecture

Our model’s architecture is designed specifically for analyzing sequence data with varying lengths and comprises three main components: (1) four convolutional blocks with pooling to distill sequence lengths, (2) five dilated convolutional blocks for capturing long-distance interactions between proteins, and (3) a cropping layer that precedes two task-specific outputs. This structure is adept at processing sequences where each element contains 480 features, utilizing max pooling to refine sequence lengths and dilated convolutions to explore distant protein interactions. Padding is applied to most sequences in preprocessing. Therefore, cropping is performed to enhance data quality.

Inspired by the Enformer (Avsec *et al*. 2021a), this architecture with tailored modifications for our dataset, our model opts for dilated convolutions over transformers due to the scale of our dataset, computational efficiency, and the unordered nature of sequence data of various lengths. Furthermore, we incorporate the Receptive-Field Attention (RFA) (Zhang *et al*. 2023) convolutional block, enabling the framework to focus on significant features through spatial attention mechanisms integrated with deep learning for advanced one-dimensional sequence data processing. Dynamic attention weights are generated via average pooling and depth-wise convolution, emphasizing the most informative sequence segments. This method aligns with our purpose to identify and analyze critical features within complex, redundant biological systems.

The RFA block’s functionality can be summarized by the equation:

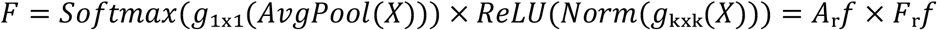

Here, X denotes the input feature map. *g*_1×1_(·) represents a depth-wise 1D convolution with a kernel size of 1 (without changing the sequence length), applied to the pooled feature map to calculate attention map *A*_r_*f. g*_*k*×*k*_(*X*) denotes a depth-wise 1D convolution on X with kernel size k, followed by batch normalization and RELU to generate transformed receptive-field spatial feature *F*_r_*f*. The final output, F, is obtained by multiplying *A*_r_*f* with *F*_r_*f*.

### Training

Due to the complexity of biological systems, particularly the multifaceted lifestyles of fungi, which often exhibit multiple forms simultaneously apart from the primary one, we utilized label smoothing to temper the confidence in our labels. label smoothing adjusts the target label (representing the true class) by reducing it by a factor of 1 − ε and distributing a small fraction ε/*C* across all classes. This results in a new, smoothed label vector:

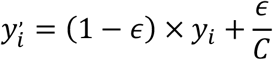

Two models were implemented using PyTorch and trained on NVIDIA RTX 4090 GPU. With a batch size set to 32 and an initial learning rate of 0.0005, we employed a learning scheduler that reduces the learning rate by 0.1 every 30 steps to enhance the stability of the training process. AdamW stochastic optimization method with weight decay as 0.1 is applied to prevent overfitting.

BPNet is trained on the same dataset using a similar strategy with the best learning rate being 0.001. Both models exhibited stabilization in their training loss and weighted F1 score around the 100th epoch. However, to ensure optimal performance, the training continued for an additional 150 epochs. The final models were selected based on their performance, specifically their weighted F1 score and AUROC on the validation set. The optimal learning rate for both models was determined through a grid search.

### Model Interpretation

For the interpretation of the model, particularly its ability to identify significant proteins, we utilized Captum to compute integrated gradients. Proteins exhibiting an attribution score greater than 0.05 were selected for further analysis. GO analysis was carried out using the pannzer2 (Törönen and Holm 2022), with the enrichment process extending to level two terms for all the proteins, facilitated by the Python package goatools.

PPI for the 59 proteins with the highest attribution values in task one and two respectively in *Verticillium alfalfae* were acquired via the STRING database (Szklarczyk *et al*. 2019) and subsequently processed using R for a detailed examination of the interaction networks.

